# Phosphite Production by *Streptomyces viridochromogenes*

**DOI:** 10.1101/2025.05.01.651723

**Authors:** Clara A. Bailey, Emma S. Broening, Brandon L. Greene

## Abstract

The industrial production of phosphochemicals is highly energy-intensive, involving the reduction of the phosphate mineral apatite to white phosphorus, a toxic and reactive intermediate with significant environmental risk. Phosphite, an activated form of phosphorus, is a potential alternative substrate for phosphochemical synthesis, yet direct reduction of phosphate to phosphite remains challenging. While environmental and microbial studies have suggested biochemical pathways for reducing phosphate to phosphite, these pathways have not been conclusively demonstrated in axenic culture. In this study, we characterize phosphite production by *Streptomyces viridochromogenes* and demonstrate that phosphite is an abiological product of phosphonoformyl-CMP decomposition, an intermediate in the biosynthesis of the herbicide phosphinothricin. The phosphonoformyl-CMP intermediate yields an “activated” phosphonoformate for decarboxylation, producing phosphite at biological temperatures and pH following phosphoanhydride hydrolysis. Using *S. viridochromogenes* spent media, we demonstrate a hybrid biotic-abiotic synthesis of the metal chelator aminotris(methylenephosphonate), illustrating a potential synthetic route to phosphochemicals from biogenic phosphite.

## INTRODUCTION

The phosphorus (P) industry lacks sustainable methods to produce phosphochemicals used in a wide range of applications, including commodity chemicals, pharmaceuticals, herbicides, flame retardants, and battery electrolytes.^1,2^ The legacy “carbothermal” process for activating P in phosphochemical synthesis reduces phosphate (PO_4_^3-^, P_i_) from apatite rock to white phosphorus (P_4_) through high-temperature processing (>1,000°C) and the use of fossil fuels as a reductant.^3^ The resulting P_4_ can be transformed into various reactive intermediates via exothermic reactions such as disproportionation to generate phosphine (PH_3_) and hypophosphorous acid (H_3_PO_2_), oxidation to form phosphorus oxides, or halogenation to form versatile phosphochemical precursors. This phosphochemical process is energy-intensive, inefficient, and produces a variety of reactive and toxic intermediates, underscoring the need for sustainable alternatives in phosphochemical production.^2^

One such alternative route to diverse phosphochemicals is the direct reduction of P_i_ to phosphite (Phi). Phi can undergo facile manipulation through Grignard-, Arbuzov- and Michael-type reactions to yield phosphonates or can disproportionate to generate PH_3_ at moderate temperatures._2,3_ Currently, Phi is produced by the carbothermal process through the oxidation of P_4_ under O_2_-limited conditions or the hydrolysis of PCl_3_. This circuitous synthetic method is employed due to the lack of viable chemical approaches to the direct two-electron reduction of P_i_ to Phi. The direct reduction of P_i_ is thermodynamically challenging with a reduction potential 280 mV more negative than proton reduction in water.^4^ Mechanochemical reduction of phosphate anhydrides, using potent hydride donors such as potassium hydride, can generate Phi; however, this process produces a complex product mixture and modest phosphite yields.^5^ Thus, developing alternative methods for selective Phi production at ambient temperatures in aqueous solution is an unsolved challenge.

**Scheme 1.**
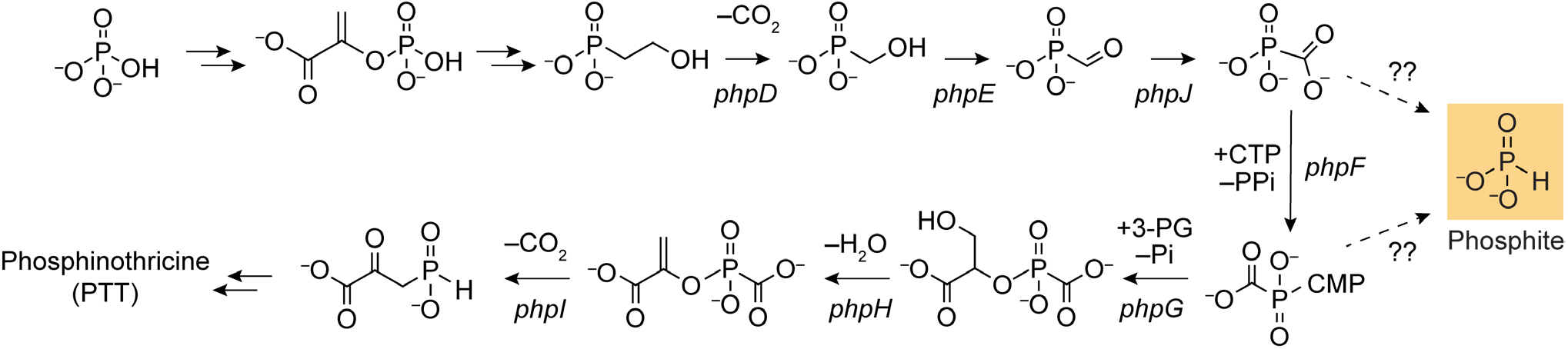
Biosynthesis pathway of bialaphos, the bioactive element of PTT.

There is substantial evidence of environmental accumulation of reduced P species, including Phi, that appear to be of biogenic origin.^6–11^ Microbial reduction of P_i_ to Phi could provide a sustainable pathway for Phi production. The microbial production of phosphonates from phosphoenolpyruvate via the enzyme PEP mutase (PepM) is well-documented.^12^ Phosphonates are an attractive “activated” P precursor in a potential microbial P_i_ reduction pathway. Yet, the conversion of phosphonates to Phi has not been conclusively demonstrated in axenic culture and, hence, the biochemical mechanism(s) involved remain largely unexplored.

In this study, we describe the Phi-producing activity of *Streptomyces viridochromogenes* via the biosynthetic pathway of the broad-spectrum herbicide and antibiotic phosphinothricin tripeptide (PTT) (**Scheme 1**). Previous study of the PTT biosynthetic pathway proposed Phi as a decarboxylation product of the intermediate phosphonoformate (PF), yet this proposal is inconsistent with the known stability of PF towards decarboxylation at near neutral pH.^13–15^ Through PF feeding experiments, knock-outs, and analysis of recombinantly expressed CTP-phosphonoformate nucleotidyltransferase (PhpF) *in vitro*, we show that Phi is produced, at biologically relevant pH and temperatures, via a phosphonoformyl-CMP intermediate. Finally, we explore the use of microbially produced Phi as a route to phosphochemical production by synthesizing the metal-chelating agent aminotris(methylenephosphonic acid) (ATMP) using spent culture medium.

## MATERIALS AND METHODS

### Material

Sodium phosphonoformate tribasic pentahydrate (PF), NaCl, KCl, 3-(N-morpholino)propanesulfonic acid (MOPS), 2-(*N*-morpholino)ethanesulfonic acid (MES), aminotris(methylenephosphonic acid) (ATMP), cytidine 5′-triphosphate disodium salt (CTP), adenosine 5′-triphosphate disodium salt (ATP), 3-(trimethylsilyl)-1-propanesulfonic acid sodium salt (TSP), NH_4_Cl, phenazine methosulfate (PMS), β-nicotinamide adenine dinucleotide (NAD^+^), resazurin, phenylmethylsulfonyl fluoride (PMSF), dithiothreitol (DTT), Triton X-100, streptomycin sulfate, imidazole, pyrophosphatase (PPase, from baker’s yeast, 500 U/mg), glycerol, 5 mm glass beads, and Amicon® centrifugal filters were purchased from Sigma Aldrich. Tris(hydroxmethyl)aminomethane hydrochloride (Tris-HCl), sodium phosphite dibasic pentahydrate, D-(+)-glucose, LB medium (Miller, powder), malt extract, and yeast extract were purchased from Thermo Fisher Scientific. Formaldehyde (37% solution) and hydrochloric acid (36.5-38.0% solution) were purchased from VWR. Kanamycin was purchased from Genesee Scientific. Ni-NTA resin was purchased from Qiagen. Phosphite dehydrogenase (PtxD) was available from a previous study.^16^

*Streptomyces viridochromogenes* DSM 40736 was obtained from the Leibniz Institute DSMZ (Braunschweig, Germany). Mutant strains WM6547, WM6601, WM6602, WM6548, WM6546, and WM6604 (ΔphpC, ΔphpD, ΔphpE, ΔphpJ, ΔphpF, and ΔphpH, respectively) were a gracious gift from Dr. William W. Metcalf.^13^

### Growth and Sampling of *Streptomyces viridochromogenes* DSM 40736

*S. viridochromogenes* colonies were transferred to 15 mL of maltose-yeast-glucose (MYG) medium (pH 6.5) and shaken at 30 °C for 2 days. These cultures were then transferred to 100 mL of MYG (with 0.1 mM or 0.5 mM PF, where indicated) with 5 mm glass beads and shaken at 30 °C for four days. Culture supernatant was collected by removing 50 mL of culture and centrifuging at 10,000 × *g* for 10 min. As a negative control, 100 mL of 0.1 mM PF in sterile MYG (pH 6.5) was incubated similarly and sampled in parallel to the live cultures. The supernatant was decanted into fresh 50 mL centrifuge tubes, and the pellet and supernatant were frozen in liquid N_2_. The supernatant was lyophilized to a powder, then resuspended in 1 mL Milli-Q H_2_O (>17 MΩ). Prior to NMR analysis, 1 mM TSP and 10% D_2_O were added for internal reference.

### NMR Analysis

NMR analysis was done at 25 °C on a Bruker Avance NEO 500 MHz spectrometer fitted with a 5 mm CryoProbe Prodigy BBO probe and acquired using TopSpin 4.1.4 software. After sample introduction, ^1^H-NMR, ^1^H-decoupled ^31^P-NMR, and ^1^H-coupled ^31^P-NMR spectra were collected. ^1^H-NMR spectra were collected using ‘zgesgp’, which features water suppression using excitation sculpting with gradients.^17 1^H-decoupled ^31^P-NMR spectra were collected with ‘P31CPD’ featuring a 2 s recycle delay, averaging 64 scans unless otherwise specified. ^1^H-coupled ^31^P-NMR spectra were collected with ‘P31’ using a 2 s recycle delay, averaging 100 scans unless otherwise specified. The methyl ^1^H peak of TSP was used as the 0.0 ppm reference. The decoupled or coupled ^31^P-NMR spectra were indirectly referenced using the TSP signal relative to 85% H_3_PO_4_.^18^ Spectral data were processed with MestReNova software.

### Quantification of Phi

Phi concentrations were determined using a fluorometric assay coupled to the activity of phosphite dehydrogenase (PtxD), as described previously.^16^ *S. viridochromogenes* spent supernatant was supplied directly to the assay without modification. Enzymatic reactions were passed through a 3 kDa Amicon® Ultra-0.5 or 3 kDa Amicon® Ultra-15 centrifugal filter unit (EMD Millipore) at 14,000 × *g* or 5,000 × *g*, respectively, for 30 min.

### *Streptomyces viridochromogenes* ΔphpH Lysate Preparation

*S. viridochromogenes* ΔphpH deletion mutant was grown in MYG liquid medium as described above. After 4 days of incubation the cells were collected and frozen in liquid N_2_. The cells were later thawed and resuspended in lysis buffer (20 mM MES, 100 mM NaCl, 10 mM PF, 1% Triton X-100, 1 mM PMSF, and 2 mM DTT at pH 6.5) at a ratio of 4 mL buffer per g cell pellet. Cells were lysed via ultrasonication (12 min processing time with a ¼” probe, 10 s on-10 s off, at 60% amplitude) and insoluble cellular material was removed by centrifugation at 14,000 × *g* for 10 min and discarded. The lysate was split into three 12.5 mL fractions, to which 5 mM ATP (from a 100 mM stock, pH 6.5) and 10 U PPase, 5 mM CTP (from a 100 mM stock, pH 6.5) and 10 U PPase, or 10 U PPase were added. The lysates were shaken at 200 rpm at 37 °C for 24 h. Afterwards, the lysates were passed through a 3 kDa centrifugal filter at 5,000 × *g* and the flow-through was collected. Of the 12 mL collected, 2 mL were reserved for Phi quantification and the remaining 10 mL were frozen in liquid N_2_ and lyophilized to a powder. For ^31^P-NMR analysis, the powder was resuspended in 0.5 mL Milli-Q H_2_O and supplied with 1 mM TSP and 10% D_2_O. Prior to use in the phosphite assay, cell lysates were diluted 1.5-fold to minimize medium effects.

### Expression and Purification of His-SUMO-PhpF

The pET-His-SUMO-PhpF plasmid was constructed using HiFi Assembly. PCR with Phusion High-Fidelity DNA Polymerase (New England Biolabs, USA) was used to generate overlapping ends for His-Smt3-NpuDnaEN102-CBD^19^ backbone and the *phpF* gene fragment. PCR products were digested with DPnI (New England Biolabs, USA) overnight at 37°C. Fragments and backbones with overlapping ends were ligated using HiFi DNA Assembly Master Mix (New England Biolabs). The resulting DNA product was transformed into *E. coli* DH5α cells and the assembly of the pET-His-SUMO-PhpF plasmid was verified by Sanger DNA sequencing.

*E. coli* BL21(DE3) cells were transformed with the pET-His-SUMO-PhpF plasmid and were cultured overnight in 10 mL LB media with 25 μg/mL kanamycin. Saturated cultures were diluted at a 1:100 ratio in fresh LB with 25 μg/mL kanamycin, grown to an OD_600_ of 0.6-0.9, induced with 0.1 mM IPTG, and incubated at 23°C for 20 h. The resulting cell pellet was resuspended in Buffer A (20 mM Tris buffer, 500 mM NaCl, and 10% v/v glycerol at pH 7.6) with 4 mL buffer per g cell pellet. The cell pellet was homogenized and lysed using a pre-cooled Avestin C3 French Press at 14,000 psi in one pass. The resulting lysate was centrifuged at 14,000 × *g* for 10 min. DNA was precipitated from the supernatant by the addition of 0.2 volume equivalents of 6% w/v streptomycin sulfate, followed by centrifugation at 14,000 × *g* for 30 min. The supernatant was then loaded on a Ni-NTA column pre-equilibrated with Buffer A. The column was washed with 30 column volumes (CVs) of Buffer B (Buffer A supplemented with 50 mM imidazole), then the desired protein was eluted with 3 CVs of Buffer C (Buffer A supplemented with 500 mM imidazole). Protein-containing fractions were combined and concentrated in a 10 kDa Amicon® Ultra-15 centrifugal filter unit at 5,000 × *g* for 10-30 min. The protein concentrate was buffer exchanged with Buffer D (20 mM MOPS, 100 mM KCl, and 10% v/v glycerol at pH 7.6) using a HiTrap desalting 5 mL column containing Sephadex G-25 resin. The resulting sample was assessed for purity using SDS-PAGE (**Supplemental Information Figure S1**).

Enzyme activity was determined using a pyrophosphate assay kit (Sigma Aldrich). Enzymatic reactions were prepared in 1.2 mL reactions with 50 mM MES (pH 6), 10 mM MgCl_2_, 5 mM PF, 15 mM CTP, and 1 μg His-SUMO-phpF. Aliquots were sampled throughout incubation at 37°C by passing 200 μL of the reaction mix through a 10 kDa Amicon® Ultra-15 centrifugal filter unit at 14,000 × *g* for 20 s. The filtrate was diluted 1:100 before use in the pyrophosphate assay (**Supplemental Information Figure S2**).

### His-SUMO-phpF Enzymatic Assays

Native activity of phpF was verified in 2 mL assay reactions containing 50 mM HEPES (pH 7.25), 10 mM MgCl_2_, 1 mM CTP, and 1 mM PF as reported previously._13_ To initiate the reaction, 5 μg of His-SUMO-phpF and 5 U PPase were added. The reaction was incubated at 37°C for 1 h before analysis by _31_P-NMR. To observe Phi-producing reactivity, 4 mL reaction mixes were prepared with 50 mM MES (pH 6 or 6.5), 10 mM MgCl_2_, 5 mM PF, 15 mM CTP, 20 μg His-SUMO-phpF, and 10 U PPase, and were incubated at 37°C. Aliquots were removed throughout the reaction for analysis with NMR and/or the fluorescence-based Phi assay. To determine if conversion to Phi was dependent on the presence of the active enzyme, a 1 mL reaction mix was prepared with 50 mM MES (pH 6), 10 mM MgCl_2_, 5 mM PF, 15 mM CTP, 5 μg His-SUMO-phpF, and 2.5 U PPase. The reaction was allowed to proceed for 2 h, after which 1 μL of trypsin (0.5 mg/mL, Promega) was added and 0.5 mL of the reaction mix was removed and immediately stored at –20°C. The remaining 0.5 mL of the reaction was incubated at 37°C for 24 h. Both trypsin-containing samples were then analyzed with NMR.

### ATMP Synthesis

The synthesis of ATMP was adapted from reported methods.^20^ *S. viridochromogenes* cultures were grown as previously described, but with the addition of 5 mM PF to culture medium and spent media collection after 4 days of growth. Spent medium (∼1 L) was collected by centrifugation at 8,000 × *g* for 30 min to remove all insoluble material. The supernatant was collected in 50 mL Falcon tubes, lyophilized to a powder, and resuspended in 10 mL Milli-Q H_2_O. The resulting 10 mL was placed in a three-necked round bottom flask with 10 mL concentrated (37%) HCl and 1.78 g NH_4_Cl. The reaction mixture was brought to reflux temperature, then 16 mL of formaldehyde was added dropwise over the course of 1 h. The reaction was refluxed for an additional 2 h, then was left at room temperature overnight. The products were precipitated by adding 150 mL acetone dropwise. The precipitate was vacuum filtered and resuspended in 15 mL of H_2_O. This crude product (710 mg) was verified with ^31^P-NMR. To increase the signal-to-noise ratio, the apodization exponential function was adjusted from 1 Hz to 4 Hz.

## RESULTS AND DISCUSSION

### Phi Production by *S. viridochromogenes*

Previous studies on the Δ*phpF* strain of *S. viridochromogenes*, grown on MYG media, suggested that Phi was produced via the accumulation of the PhpF substrate, PF, and its spontaneous decarboxylation to Phi.^13^ To replicate and potentially enhance Phi production, we cultivated *S. viridochromogenes* in MYG medium supplemented with 500 μM PF.^13^ After 4 days of growth, we observed a putative Phi peak at 2.91 ppm by ^31^P-NMR (**Figure 1A**). The identity of the Phi peak is supported by ^1^H-coupled ^31^P-NMR, with a characteristically large ^1^H coupling constants (725 Hz), indicative of a direct P-H bond, and by comparison with an authentic Phi standard (**Supplemental Information Figure S3**).^21^ Minor differences in chemical shift and ^1^H-^31^P coupling strength between the Phi signal from *S. viridochromogenes* media and the authentic standard are expected and reasonable based on the medium-dependence of the ionizable Phi.^22– 24^ We further confirmed the identity of the ^31^P-NMR signal by spiking Phi and P_i_ into the spent media sample, demonstrating that the peak was indeed Phi and not attributable to inorganic P_i_ (**Supplemental Information Figure S4**). Using our previously developed enzyme-coupled fluorometric assay, we also quantified the Phi product.^16^ Following the 4 day growth, the cell culture medium contained 110 μM ± 40 μM Phi (**Figure 1B**), indicating 22% conversion of PF to Phi in culture. When PF was incubated in sterile MYG medium, no Phi was observed, either by ^31^P-NMR or the enzymatic Phi assay (**Supplemental Information Figure S5**). These results confirm that PF does not spontaneously decompose to Phi over the time scale, temperature, and pH of the *S. viridochromogenes* cultures.^15^

**Figure 1.**
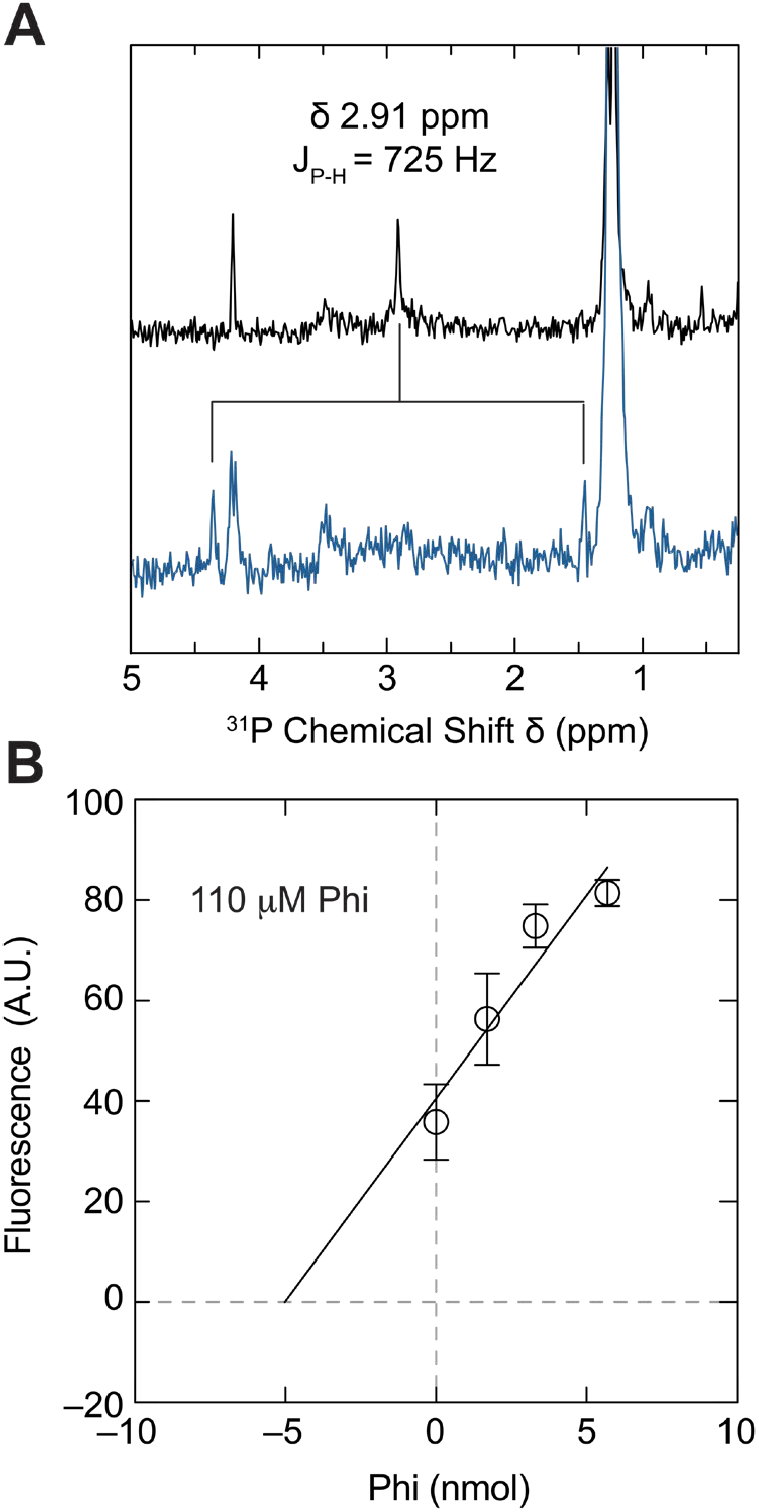
Phi signal in *S. viridochromogenes* spent medium. **A** Proton-decoupled (black) and proton-coupled (blue) ^31^P-NMR spectra of *S. viridochromogenes* spent medium after 4 days of incubation at 30°C in MYG with 500 μM PF, concentrated 100-fold. Splitting of the Phi peak at 2.91 ppm is shown along with the coupling constant, J_P-H_. **B** Quantification of Phi in *S. viridochromogenes* unconcentrated spent medium using the Phi assay. The standard addition curve reading was 110 ± 40 μM Phi.

We observe a Phi signal in the culture supernatant as early as 48 h after inoculation, which persisted through the remaining 2 days of culturing (**Supplemental Information Figure S6**). At the same 48 h time point, we also detected other phosphonate intermediates of the PTT pathway in the spent medium supernatant. To determine the dependence of the Phi signal on supplemented PF, we grew *S. viridochromogenes* cultures with and without 500

μM PF in MYG medium. In all biological replicates, we detected Phi only in cultures supplied with PF. The supplementation of PF also increased the quantity of several phosphonate intermediates in the PTT pathway (δ 15-45 ppm, **Supplemental Information Figure S7**), suggesting that the PTT biosynthetic genes are expressed and that PTT production is regulated via the production of a PF precursor.^25^ This correlation between Phi and phosphonate production implicates the PTT pathway in Phi biosynthesis but does not establish molecular specificity.

### *S. viridochromogenes* Deletion Mutants

We analyzed the spent media of *S. viridochromogenes* mutants with genetic deletions in the PTT biosynthetic pathway (Δ*phpC*, Δ*phpD*, Δ*phpE*, Δ*phpJ*, Δ*phpF*, and Δ*phpH*), supplemented with 500 μM PF, to determine if any of these genes contributed to Phi production. As expected, deletions in the *phn* operon of genes involved in the biosynthesis of the PF intermediate (Δ*phpC*, Δ*phpD*, Δ*phpE*, and Δ*phpJ*, **Scheme 1**) did not prevent the production of PTT (δ 42 ppm) from exogenously supplied PF. Conversely, deletions of genes downstream from PF synthesis (Δ*phpF* and Δ*phpH*) showed no PTT signal (**Supplemental Information Figure S8**). Unexpectedly, all deletion mutants, except for Δ*phpF*, show Phi signal (**Figure 2**). Previous studies on the Δ*phpF S. viridochromogenes* mutant identified a peak by ^31^P-NMR that was attributed to Phi. We also observed a ^31^P-NMR peak in Δ*phpF* growth medium supernatant in the vicinity of Phi signals (δ 4.45 ppm); however, this species did not exhibit the characteristically large ^1^H-^31^P coupling of Phi in the ^1^H-coupled ^31^P-NMR (J_H-P_ ∼10 Hz), suggesting it is a phosphonate, but not Phi. Based on these genetic knockout studies, the molecular origins of Phi appear to be a result of PhpF activity in *S. viridochromogenes* (**Scheme 1**).

**Figure 2.**
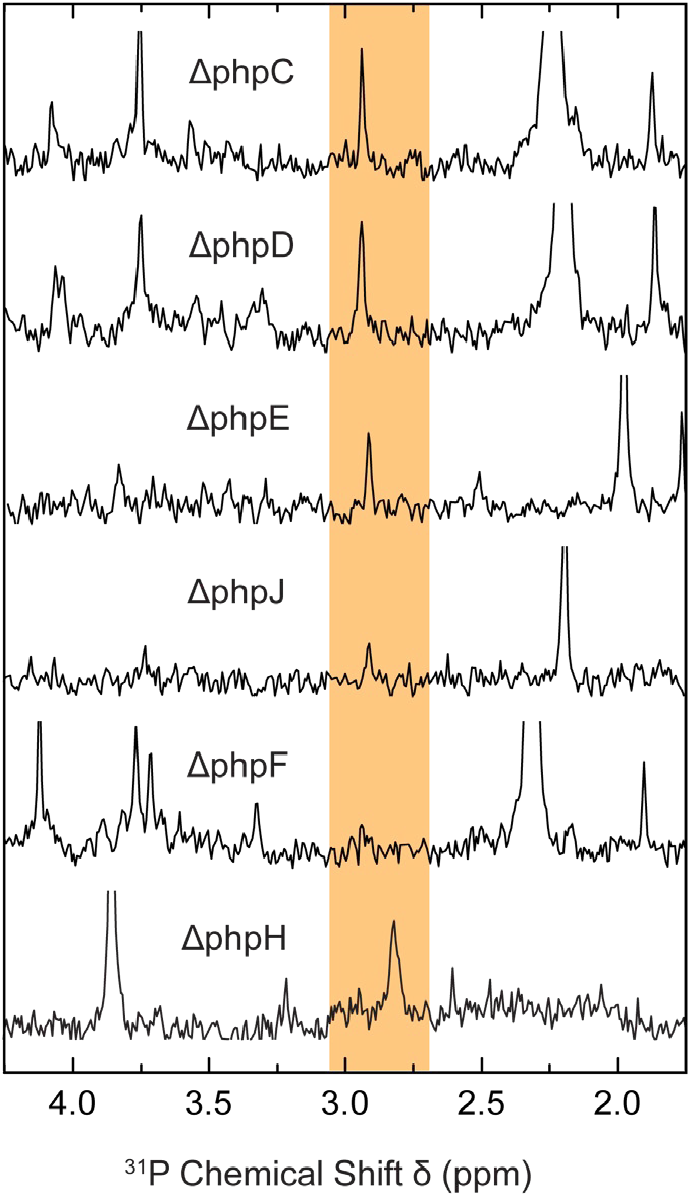
Proton-decoupled ^31^P-NMR spectrum of *S. viridochromogenes* deletion mutant spent media after 4 days of incubation at 30°C in MYG with 100 μM PF, concentrated 100-fold. Putative Phi peaks are highlighted in orange.

To separate Phi production from cellular growth and confirm the Phi signal’s dependence on enzymatic activity, we assayed the soluble lysate of *S. viridochromogenes*

Δ*phpH* for Phi production. When supplemented with 10 mM PF, the lysate produced Phi, but only in the presence of PPase and CTP (**Figure 3**). Although spectral congestion prevented the identification of Phi by ^1^H-decoupled ^31^P-NMR, ^1^H-coupled ^31^P-NMR revealed an emergent peak corresponding to the Phi down-field peak of the doublet at 4.48 ppm, as well as signals consistent with phosphonoformyl-CMP (PF-CMP, **Figure 3**).^13^ To confirm the Phi signal, we quantified the Phi concentration using the enzymatic Phi assay, which showed 73 ± 7 μM Phi in the CTP-supplemented lysate and no Phi production in the absence of PPase or CTP (**Figure 3, inset**). The proposed function of PhpF is that of a CTP-phosphonoformate nucleotidyltransferase. The concurrent production of PF-CMP and Phi suggested that PhpF is responsible for the formation of Phi via a PF-CMP intermediate.^26^

**Figure 3.**
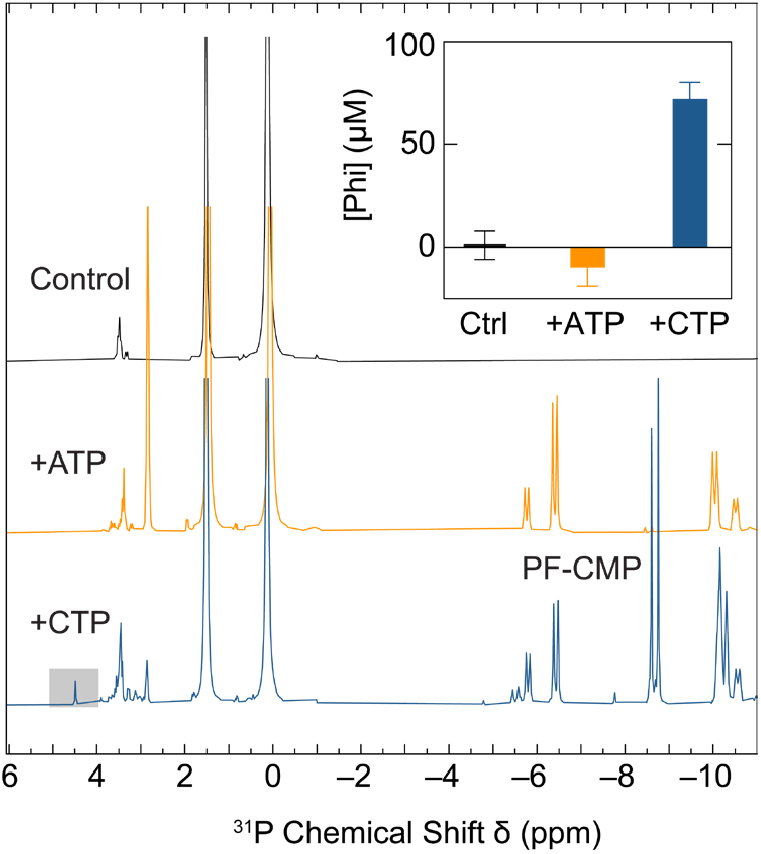
Phi production in *S. viridochromogenes* ΔphpH lysate. Proton-coupled ^31^P-NMR spectra of lysate incubated with 10 U PPase (control), 10 mM ATP and 10 U PPase, or 10 mM CTP and 10 U PPase. A putative Phi doublet peak at 4.48 ppm is highlighted. Inset: Quantification of Phi in cell lysates using the Phi assay.

**Scheme 2.**
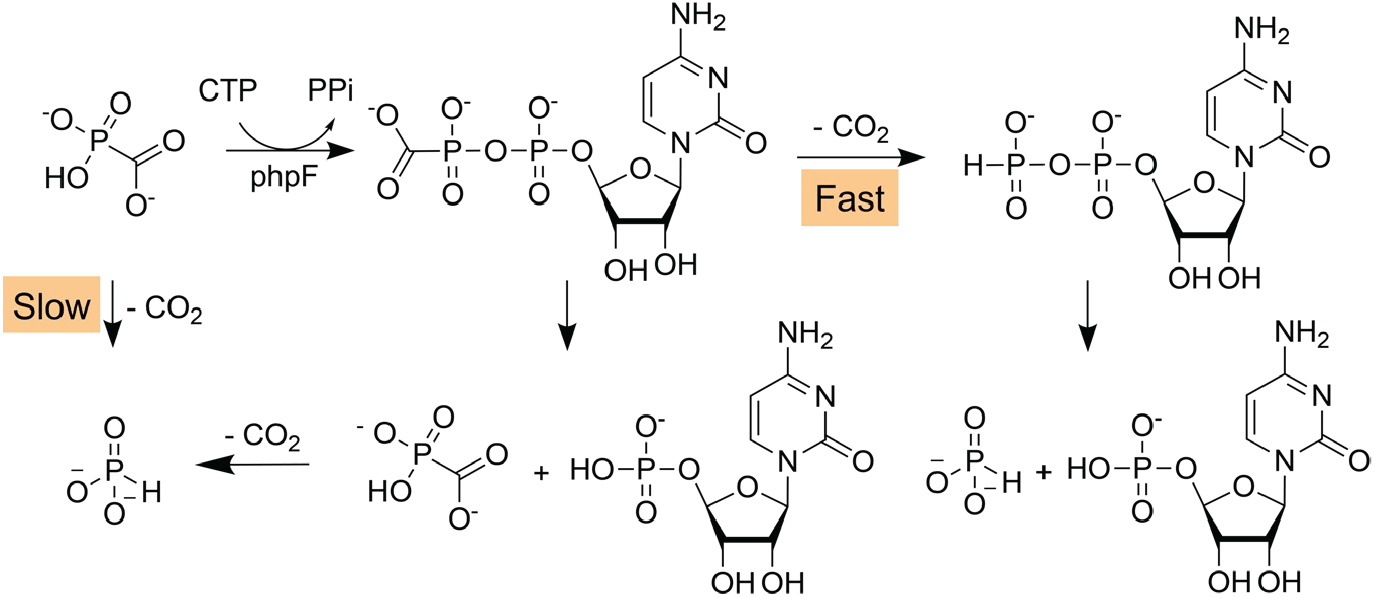
Proposed mechanism for the formation of Phi by *S. viridochromogenes*.

### Enzymatic Assays with Recombinant His-SUMO-PhpF

We hypothesized that activation of PF by PhpF promotes the decarboxylation of PF-CMP, yielding a phosphinated intermediate (Phi-CMP) that could decompose to Phi and CMP upon the hydrolysis of the phosphoanhydride bond (**Scheme 2**). To test this hypothesis, we recombinantly expressed and purified PhpF from *E. coli* for analysis. The enzyme exhibited the expected reactivity in the presence of PF, CTP, MgCl_2_, and PPase, fully converting PF to PF-CMP over 1 h at 37°C (**Supplemental Information Figure S9**). To counteract PF-CMP hydrolysis to CMP and PF, we added excess CTP to the reaction mixture, and carried out the reaction at pH 6 to favor decarboxylation by CMP-PF. After 2 h of incubation at 37°C, we observed PF-CMP, CTP, and P_i_ (**Figure 4**, top). After 24 h of incubation, we observed two new peaks, a singlet at 2.74 ppm and a doublet at 5.36 ppm with a splitting of 750 Hz and 823 Hz, respectively (**Figure 4**, middle) that were not present in the PhpF-free control after 24 h (**Figure 4**, bottom). Based on the chemical shift and J_P-H_, we assign the 2.74 ppm peak to Phi, and the 5.36 ppm doublet to Phi-CMP. To determine if the conversion of PF-CMP to Phi-CMP was enzymatic or spontaneous, we treated the reaction with trypsin after 2 h of incubation to digest PhpF. The digested sample still produced Phi from PF-CMP over 24 h (**Supplemental Information Figure S10**). We again quantified the Phi production rate using the enzymatic Phi assay (**Figure 5**). At pH 6, Phi was produced at a rate of 8.7 μM Phi/h. The relatively high activity of PhpF for its native reaction (*k*_cat_ of 60 ± 10 s^-1^) and the observation of Phi production from trypsin-digested enzyme assays (**Supplemental Information Figure S2**), implies Phi production is nonenzymatic via the decomposition of Phi-CMP. Given the prevalence of cytidyltransferase activation in phosphonate biosynthesis,^27^ these reactive intermediates may play a larger role in metabolite leakage in the environment.

**Figure 4.**
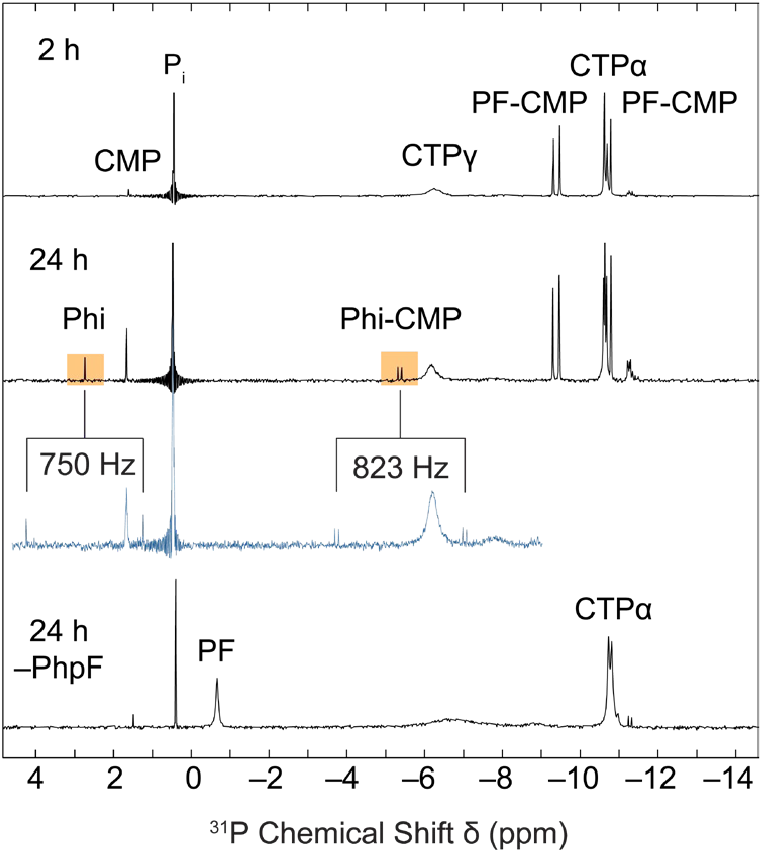
Phi production in phpF enzymatic assays. Proton-decoupled ^31^P-NMR spectra of a phpF assay reaction (50 mM MES, pH 6, 10 mM MgCl_2_, 5 mM PF, 15 mM CTP, 5 μg/mL His-SUMO-phpF, and 5 U/mL PPase) after 2 h and 24 h (black spectra, top) incubation at 37°C. The proton-coupled ^31^P-NMR spectrum of the reaction after 24 h is shown in blue. A proton-decoupled ^31^P-NMR spectrum of a control reaction without His-SUMO-phpF after 24 h incubation is shown (bottom).

**Figure 5.**
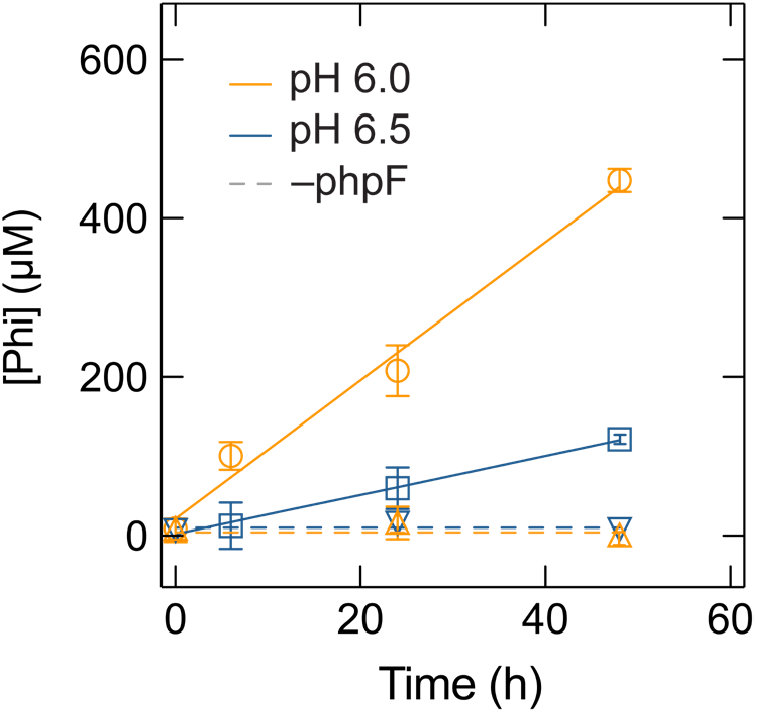
Conversion of PF to Phi by His-SUMO-phpF activity. The Phi yield with time of the phpF assay reaction is shown at pH 6 (orange) and pH 6.5 (blue). Reactions where His-SUMO-phpF was inactivated by heat before addition to the reaction mixture are shown with triangles and dashed lines.

### ATMP Synthesis

ATMP is a metal chelator with a wide range of industrial applications including the prevention of scaling, corrosion, and use in detergents.^28–30^ ATMP can be produced through a facile one-pot Mannich-type reaction involving phosphorous acid, formaldehyde, and ammonia.^20^ To demonstrate the utility of *S. viridochromogenes* in P activation, we used Phi-rich spent media as a feedstock for a hybrid biological-abiological synthesis of ATMP. To optimize phosphonate synthesis, and consequently Phi production by *S. viridochromogenes*, we grew cultures with an order of magnitude greater PF concentration in the medium (5 mM). The resulting concentrated spent media was able to provide phosphorous acid in the ATMP synthesis reaction, resulting in a proton-decoupled ^31^P-NMR peak at 7.43 ppm (**Figure 6**, top). This is in good agreement with literature values of 7.8 ppm^31^ and our own analysis of an ATMP standard (7.26 ppm, **Supplementary Figure S11**). Proton-coupled ^31^P-NMR showed an expected set of triplet peaks due to coupling with neighboring methylene groups. While the overall yield was modest from 1 L of culture (700 mg crude product), this proof-of-concept synthesis demonstrates that *S. viridochromogenes* cultures can serve as a feedstock for the synthesis of industrially relevant reduced phosphorus products. Furthermore, optimization of the biosynthetic pathway to both PF and PF-CMP from P_i_ and integration into a more atom and energy-efficient host system is expected to drastically improve biotechnological access to diverse phosphochemicals via microbially produced Phi.

**Figure 6.**
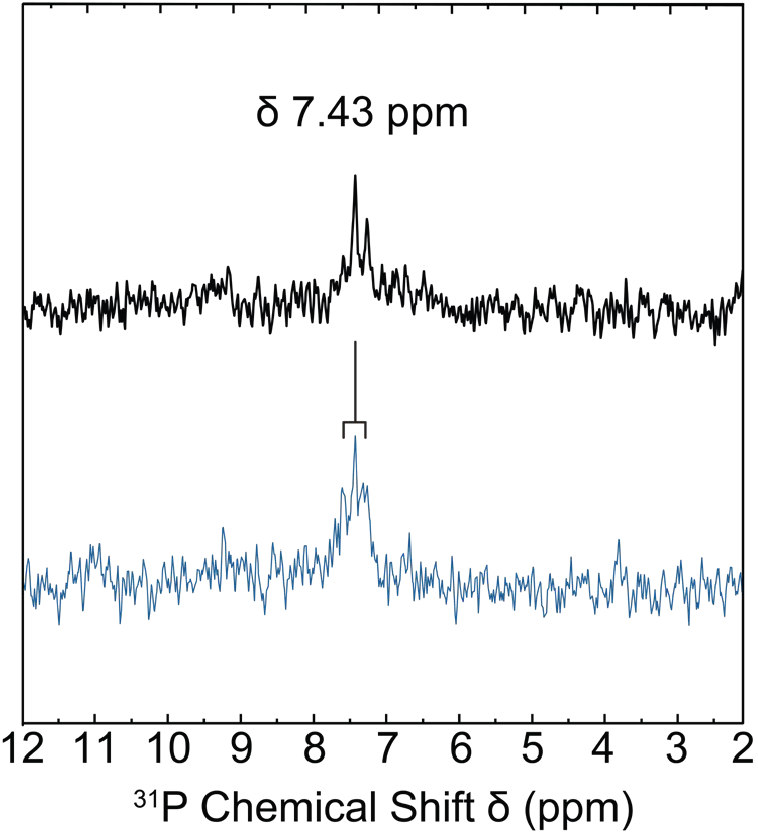
Preparation of ATMP from *S. viridochromogenes* spent medium. ATMP synthesis crude product was resuspended in 10% D_2_O and analyzed with proton-decoupled (black) and proton-coupled (blue) ^31^P-NMR. The major product and putative ATMP peak is labeled. Splitting of the ATMP peak due to neighboring methylene groups is shown.

## CONCLUSION

Microbial Phi production presents a promising alternative to energy-intensive legacy methods. Here we show Phi production in *S. viridochromogenes* as a byproduct of the PTT biosynthesis pathway through the activity of the PF-activating enzyme PhpF. In addition, we show that *S. viridochromogenes* culture can serve as a feedstock for the industrially relevant chemical synthesis of ATMP. The characterization of a Phi-producing pathway in microbial metabolism opens the potential for discovery of other microbial means of Phi generation, perhaps similarly related to phosphonate metabolism. Further work in microbial Phi production will have significant implications for sustainable reduced P reduction as well as the global biogeochemical P cycle.

## ASSOCIATED CONTENT

### Supporting Information

The Supporting Information is available free of charge on the ACS Publications website.

## AUTHOR INFORMATION

### Author Contributions

The manuscript was written through contributions of all authors. All authors have given approval to the final version of the manuscript.

## ACKNOWLEDGMENT

We gratefully acknowledge Prof. William W. Metcalf for supplying *S. viridochromogenes* deletion mutants and Dr. Hongjun Zhou for assistance with NMR experiments.

## REFERENCES

(1) Protasiewicz, J. D. From Rock-Stable to Reactive Phosphorus. Science 2018, 359, 1333–1333.

(2) Geeson, M. B.; Cummins, C. C. Let’s Make White Phosphorus Obsolete. ACS Cent. Sci. 2020, 6, 848–860.

(3) Bettermann, G.; Krause, W.; Riess, G.; Hofmann, T. Phosphorus Compounds, Inorganic. In Ullmann’s Encyclopedia of Industrial Chemistry; John Wiley & Sons, Ltd, 2000.

(4) Pourbaix, M. Atlas of Electrochemical Equilibria in Aqueous Solutions, [1st English ed.].; Pergamon Press: Oxford, 1966.

(5) Zhai, F.; Xin, T.; Geeson, M. B.; Cummins, C. C. Sustainable Production of Reduced Phosphorus Compounds: Mechanochemical Hydride Phosphorylation Using Condensed Phosphates as a Route to Phosphite. ACS Cent. Sci. 2022, 8, 332–339.

(6) Dévai, I.; Felföldy, L.; Wittner, I.; Plósz, S. Detection of Phosphine: New Aspects of the Phosphorus Cycle in the Hydrosphere. Nature 1988, 333, 343–345.

(7) Devai, I.; DeLaune, R. D.; Devai, G.; W.H. Patrick, Jr.; Czegeny, Phosphine Production Potential of Various Wastewater and Sewage Sludge Sources. Anal. Lett. 1999, 32, 1447–1457.

(8) McDowell, M. M.; Ivey, M. M.; Lee, M. E.; Firpo, V. V. V. D.; Salmassi, T. M.; Khachikian, C. S.; Foster, K. L. Detection of Hypophosphite, Phosphite, and Orthophosphate in Natural Geothermal Water by Ion Chromatography. J. Chromatogr. A 2004, 1039, 105–111.

(9) Han, C.; Geng, J.; Xie, X.; Wang, X.; Ren, H.; Gao, S. Determination of Phosphite in a Eutrophic Freshwater Lake by Suppressed Conductivity Ion Chromatography. Environ. Sci. Technol. 2012, 46, 10667–10674.

(10) Van Mooy, B. A. S.; Krupke, A.; Dyhrman, S. T.; Fredricks, H. F.; Frischkorn, K. R.; Ossolinski, J. E.; Repeta, D. J.; Rouco, M.; Seewald, J. D.; Sylva, S. P. Major Role of Planktonic Phosphate Reduction in the Marine Phosphorus Redox Cycle. Science 2015, 348, 783–785.

(11) Repeta, D. J.; Ferrón, S.; Sosa, O. A.; Johnson, C. G.; Repeta, L. D.; Acker, M.; DeLong, E. F.; Karl, D. M. Marine Methane Paradox Explained by Bacterial Degradation of Dissolved Organic Matter. Nat. Geosci. 2016, 9, 884–887.

(12) Horsman, G. P.; Zechel, D. L. Phosphonate Biochemistry. Chem. Rev. 2017, 117, 5704–5783.

(13) Blodgett, J. A. V.; Thomas, P. M.; Li, G.; Velasquez, J. E.; van der Donk, W. A.; Kelleher, N. L.; Metcalf, W. W. Unusual Transformations in the Biosynthesis of the Antibiotic Phosphinothricin Tripeptide. Nat. Chem. Biol. 2007, 3, 480–485.

(14) Warren, S.; R. Williams, M. The Acid-Catalysed Decarboxylation of Phosphonoformic Acid. J. Chem. Soc. B: Phys. Org. 1971, 0, 618–621.

(15) Bundgaard, H.; Mork, N. Kinetics of the Decarboxylation of Foscarnet in Acidic Aqueous Solution and Its Implication in Its Oral Absorption. Int. J. Pharm. 1990, 63, 213–218.

(16) Bailey, C. A.; Greene, B. L. A Fluorometric Assay for High-Throughput Phosphite Quantitation in Biological and Environmental Matrices. Analyst 2023, 148, 3650–3658.

(17) Hwang, T. L.; Shaka, A. J. Water Suppression That Works. Excitation Sculpting Using Arbitrary Wave-Forms and Pulsed-Field Gradients. J. Mag. Res. A 1995, 112, 275–279.

(18) Harris, R. K.; Becker, E. D.; Cabral de Menezes, S. M.; Granger, P.; Hoffman, R. E.; Zilm, K. W. Further Conventions for NMR Shielding and Chemical Shifts (IUPAC Recommendations 2008). Pure App. Chem. 2008, 80, 59–84.

(19) Guerrero, F.; Ciragan, A.; Iwaï, H. Tandem SUMO Fusion Vectors for Improving Soluble Protein Expression and Puriication. Protein Expres. Purif. 2015, 116, 42–49.

(20) Moedritzer, K.; Irani, R. R. The Direct Synthesis of α-Aminomethylphosphonic Acids. Mannich-Type Reactions with Orthophosphorous Acid. J. Org. Chem. 1966, 31, 1603–1607.

(21) Majumdar, A.; Sun, Y.; Shah, M.; Freel Meyers, C. L. Versatile ^1^ H− ^31^ P− ^31^ P COSY 2D NMR Techniques for the Characterization of Polyphosphorylated Small Molecules. J. Org. Chem. 2010, 75 (10), 3214–3223.

(22) Smithers, G. W.; O’Sullivan, W. J. 31P Nuclear Magnetic Resonance Study of Phosphoribosyldiphosphate and Its Interaction with Magnesium Ions. J. Biol. Chem. 1982, 257, 6164–6170.

(23) Eykyn, T. R.; Kuchel, P. W. Scalar Couplings as pH Probes in Compartmentalized Biological Systems: ^31^ P NMR of Phosphite. Magn. Reson. Med. 2003, 50, 693–696.

(24) Moedritzer, K. pH Dependence of Phosphorus-31 Chemical Shifts and Coupling Constants of Some Oxyacids of Phosphorus. Inorg. Chem. 1967, 6, 936–939.

(25) Ju, K.-S.; Nair, S. K. Convergent and Divergent Biosynthetic Strategies towards Phosphonic Acid Natural Products. Curr. Opin. Chem. Biol. 2022, 71, 102214.

(26) Blodgett, J. A.; Zhang, J. K.; Yu, X.; Metcalf, W. W. Conserved Biosynthetic Pathways for Phosalacine, Bialaphos and Newly Discovered Phosphonic Acid Natural Products. J. Antibiot. 2016, 69, 15–25.

(27) Rice, K.; Batul, K.; Whiteside, J.; Kelso, J.; Papinski, M.; Schmidt, E.; Pratasouskaya, A.; Wang, D.; Sullivan, R.; Bartlett, C.; Weadge, J. T.; Van Der Kamp, M. W.; Moreno-Hagelsieb, G.; Suits, M. D.; Horsman, G. P. The Predominance of Nucleotidyl Activation in Bacterial Phosphonate Biosynthesis. Nat. Commun. 2019, 10, 3698.

(28) Bordas, F.; Bourg, A. C. M. Effect of Complexing Agents (EDTA and ATMP) on the Remobilization of Heavy Metals from a Polluted River Sediment. Aquat. Geochem. 1998, 4, 201–214.

(29) Armbruster, D.; Müller, U.; Happel, O. Characterization of Phosphonate-Based Antiscalants Used in Drinking Water Treatment Plants by Anion-Exchange Chromatography Coupled to Electrospray Ionization Time-of-Flight Mass Spectrometry and Inductively Coupled Plasma Mass Spectrometry. J. Chromatogr. A 2019, 1601, 189–204.

(30) Jaworska, J.; Van Genderen-Takken, H.; Hanstveit, A.; Van De Plassche, E.; Feijtel, T. Environmental Risk Assessment of Phosphonates, Used in Domestic Laundry and Cleaning Agents in the Netherlands. Chemosphere 2002, 47, 655–665.

(31) Oromí-Farrús, M.; Minguell, J. M.; Oromi, N.; Canela-Garayoa, R. A Reliable Method for Quantiication of Phosphonates and Their Impurities by ^31^ P NMR. Anal. Lett. 2013, 46, 1910–1921.

