## Supplemental Information for "Phosphite Production by *Streptomyces viridochromogenes*"

### Table of Contents

**Figure S1.** SDS-PAGE gel describing the purification of His-SUMO-phpF S3

**Figure S2.** His-SUMO-phpF activity by assaying for pyrophosphate S4

**Figure S3.** $^{31}$P-NMR spectrum of Phi standard S5

**Figure S4.** Phi signal confirmation with Phi- and Pi-spiking S6

**Figure S5.** PF stability in sterile MYG medium S7

**Figure S6.** $^{31}$P-NMR spectrum of *S. viridochromogenes* spent media sampled throughout incubation S8

**Figure S7.** Phi signal dependence on PF in *S. viridochromogenes* spent medium S9

**Figure S8.** $^{31}$P-NMR spectra of *S. viridochromogenes* deletion mutants S10

**Figure S9.** $^{31}$P-NMR spectra confirming CTP-phosphonoformate nucleotidyltransferase activity of His-SUMO-phpF S11

**Figure S10.** $^{31}$P-NMR spectra demonstrating Phi production in trypsin-digested His-SUMO-phpF reaction mixtures S12

**Figure S11.** $^{31}$P-NMR spectrum of ATMP standard S13

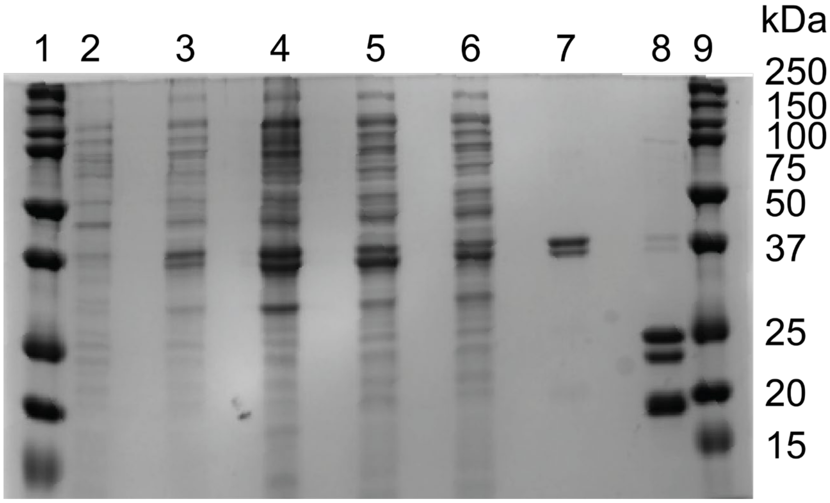


**Supplemental Information Figure S1.** SDS-PAGE gel describing the purification of His-SUMO-phpF. Lanes contain the *E. coli* BL21(DE3) pellet prior to induction (2), the *E. coli* BL21(DE3) pellet with overexpressed His-SUMO-phpF (3), the cell pellet after lysis (4), the cell lysate (5), Ni-NTA column flow-through (6), eluted final protein (7), and ULP1-digested protein product (8) and molecular weight ladders (1,9). The His-SUMO-phpF theoretical mass is 35,444 Da.

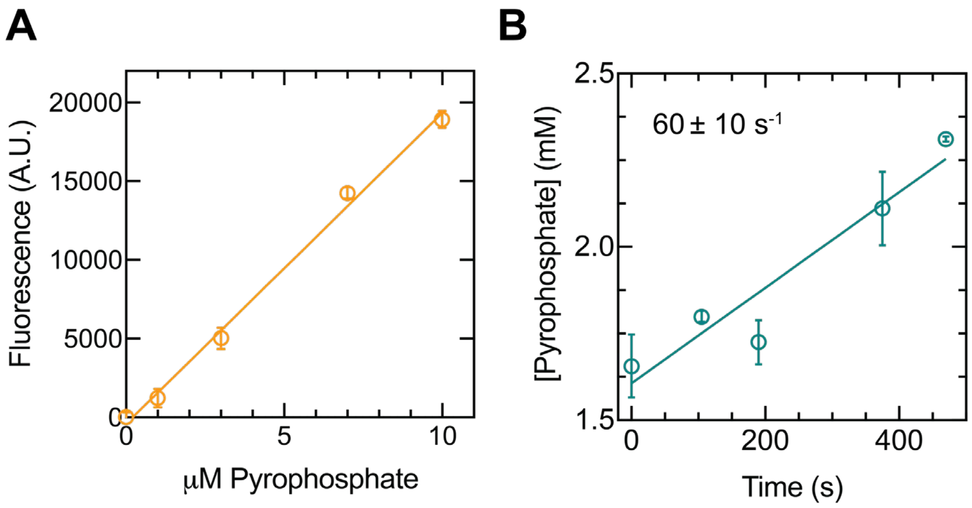


**Supplemental Information Figure S2.** Quantifying His-SUMO-phpF activity by assaying for pyrophosphate. **A** Standard curve showing fluorescence response in the linear range of the pyrophosphate assay. **B** Example kinetics experiment for the production of pyrophosphate during the formation of phosphonoformyl-CMP from phosphonoformate and CTP.

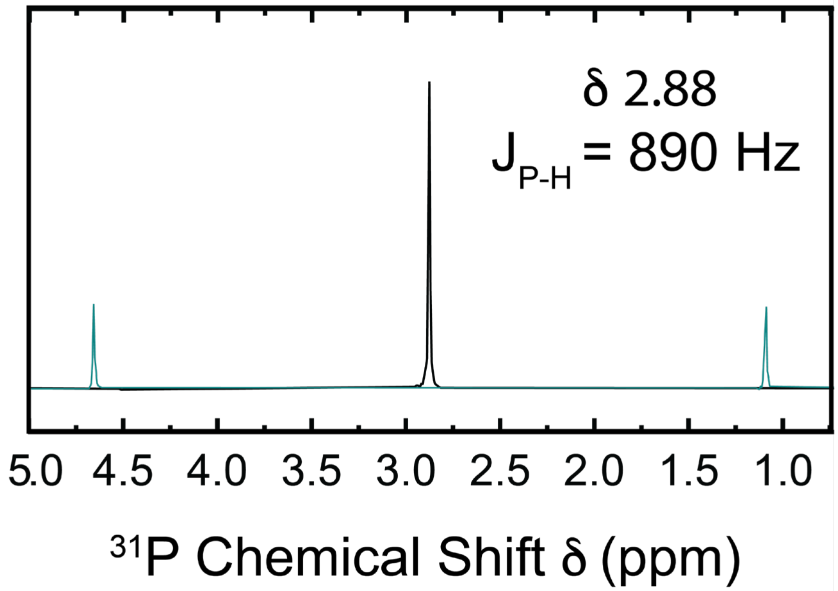


**Supplemental Information Figure S3.** $^{31}$P-NMR proton-decoupled (black) and -coupled (blue) spectra of 10 mM Phi (pH 7).

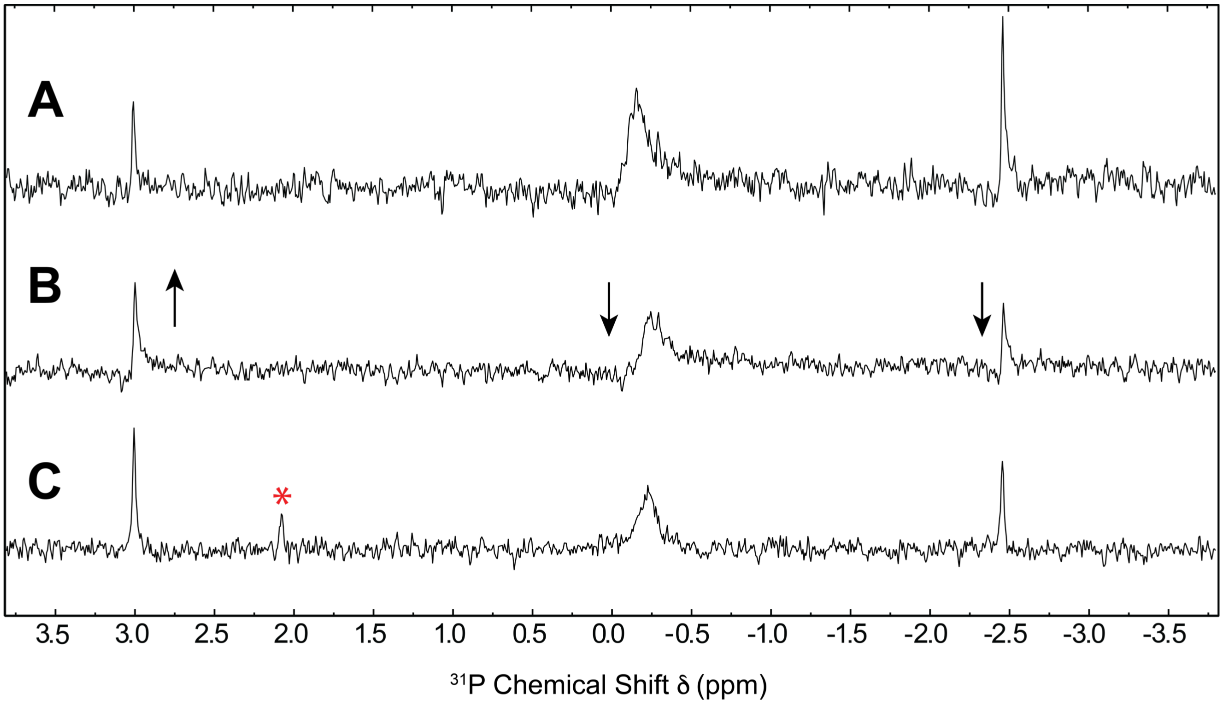


**Supplemental Figure S4.** Phi signal confirmation with Phi- and $P_i$-spiking. **A** Proton-decoupled $^{31}$P-NMR spectra of *S. viridochromogenes* spent medium after 4 days of growth in MYG with 100 µM PF, concentrated 100-fold. Spectrum was acquired with 48 scans. **B** Sample from **A** spiked with 200 µM Phi. Spectrum was acquired with 24 scans. Signals indicated with a downward-facing arrow decreased in peak area by 53-54%. Putative Phi signal indicated with an upward-facing arrow increased in peak area by 49%. **C** Sample from **B** spiked with 200 µM $P_i$. The appearance of a new peak at 2.08 ppm is indicated with an asterisk.

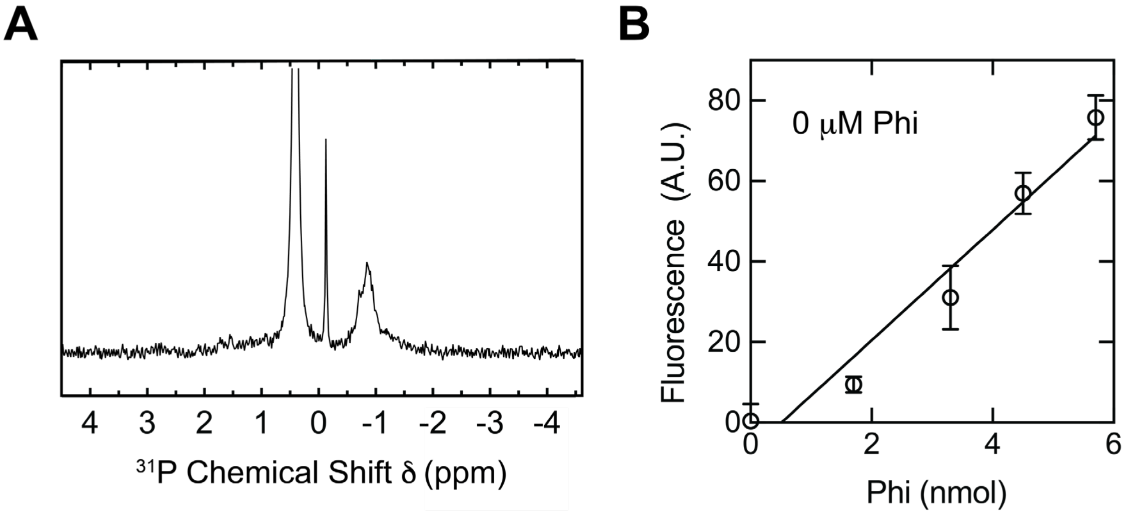


**Supplemental Information Figure S5.** PF stability in sterile MYG medium. **A** Proton-decoupled $^{31}$P-NMR spectrum of 100 μM PF in sterile MYG, incubated at 30°C for 4 days. **B** Quantification of Phi in sterile MYG supplied with PF. The standard addition curve reading was −11 ± 9 μM Phi (0 nmol).

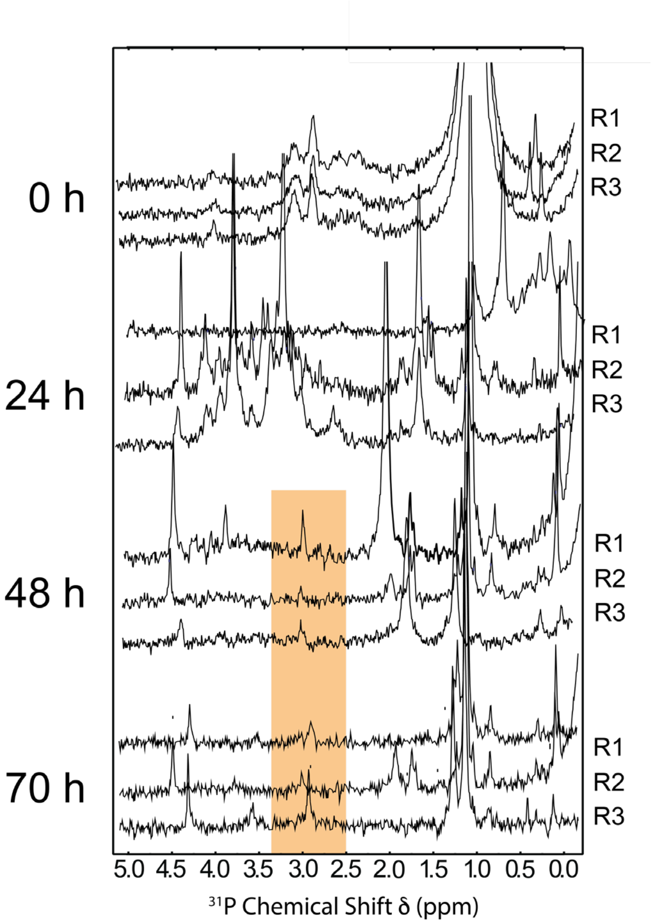


**Supplemental Figure S6.** Proton-decoupled $^{31}$P-NMR spectrum of *S. viridochromogenes* spent media sampled throughout incubation at 30°C in MYG with 500 μM PF, concentrated 100-fold. Three biological replicates (R1-R3) are shown for each timepoint. Putative Phi peaks are highlighted.

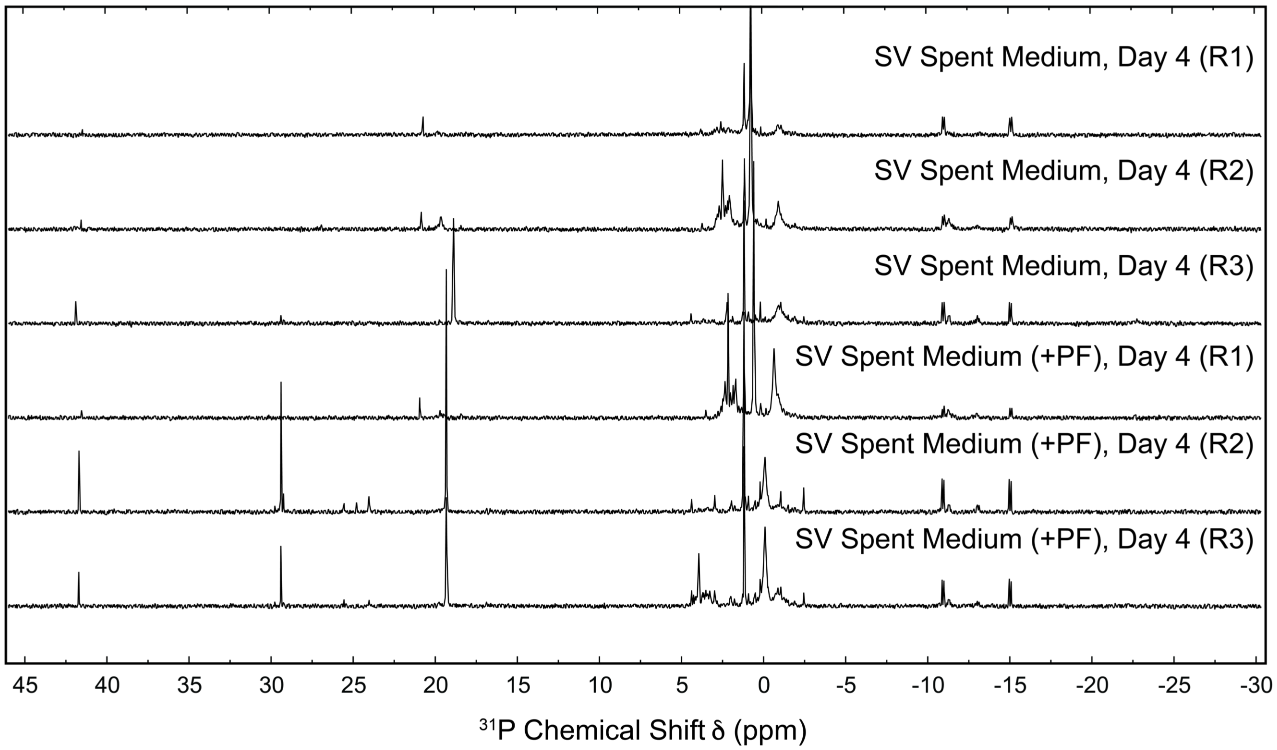


**Supplemental Figure S7.** Phi signal dependence on PF in *S. viridochromogenes* spent medium. Proton-decoupled $^{31}$P-NMR spectrum of *S. viridochromogenes* spent medium after 4 days of incubation at 30°C in MYG with (bottom) or without (top) 500 μM PF, concentrated 100-fold. Three biological replicates are shown (R1-R3).

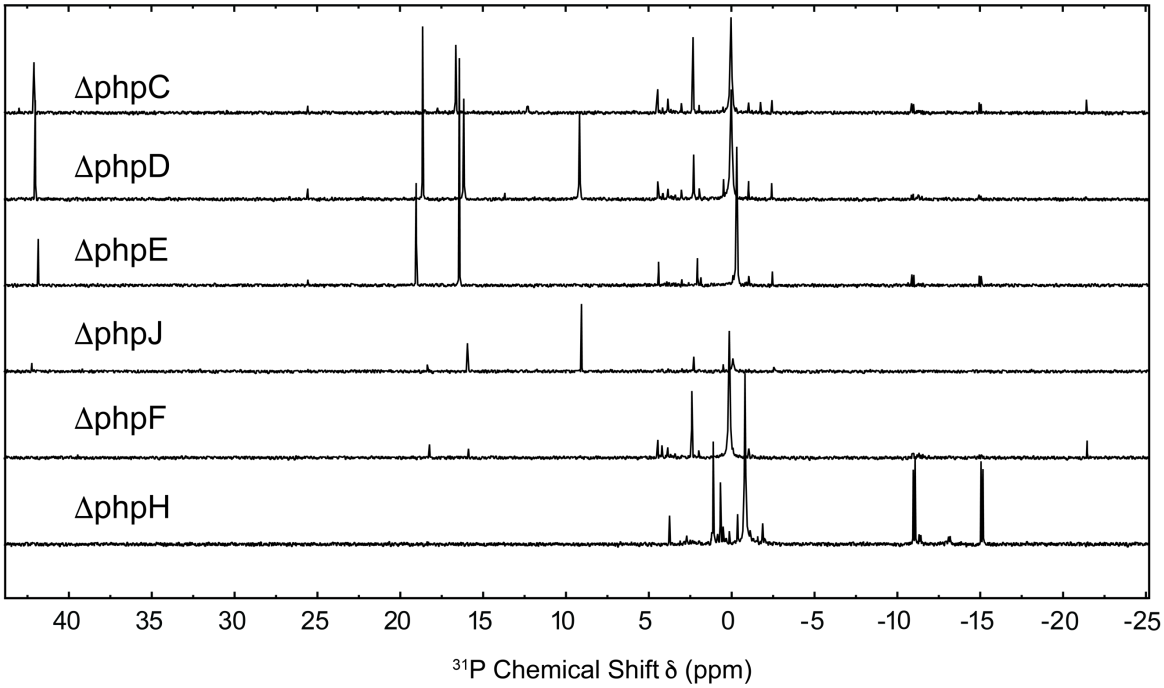


**Supplemental Figure S8.** Proton-decoupled $^{31}$P-NMR spectra of *S. viridochromogenes* deletion mutants. Cultures were grown in MYG with 500 μM PF for 4 days and the spent media was concentrated 100-fold prior to analysis. The ΔphpC mutant shows the expected accumulation of 2-aminomethylphosphonate (δ 16.7). The ΔphpE and ΔphpD mutants similarly show expected intermediate accumulation (hydroxymethyl phosphonate, δ 16.7, and hydroxyethyl phosphonate, δ 18.7 , respectively) and Phi production. The ΔphpH mutant shows no accumulation of any intermediate in the PTT biosynthesis pathway, in accordance with the results of Blodgett *et al.*

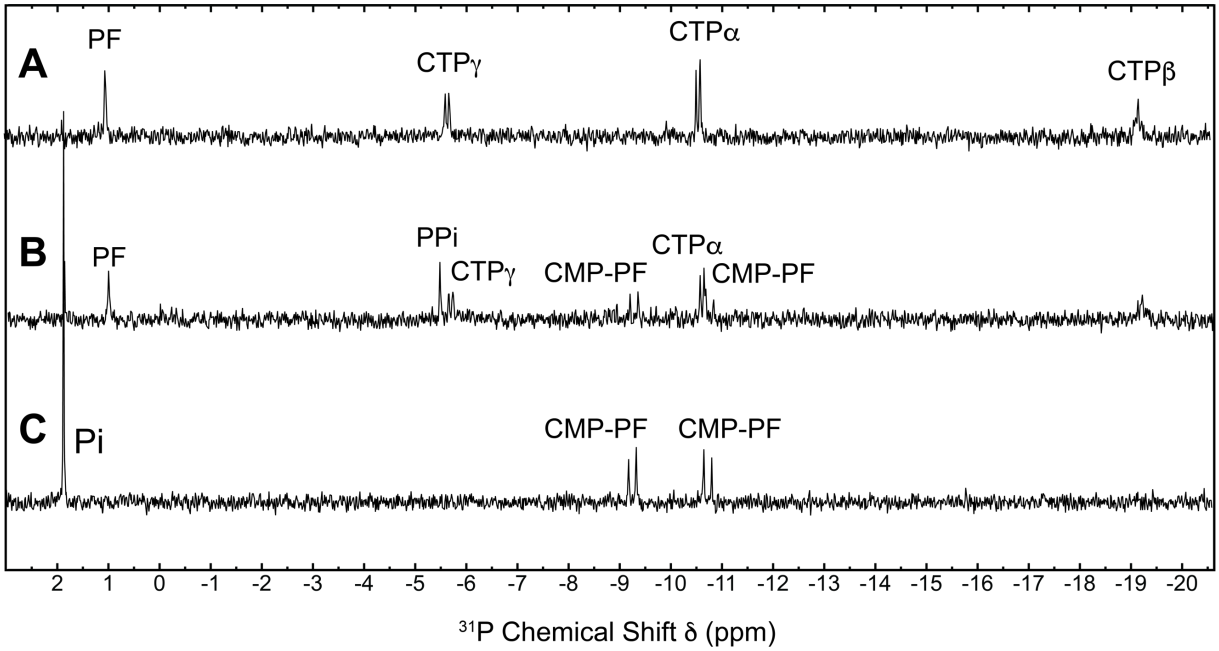


**Supplemental Figure S9.** Proton-decoupled $^{31}$P-NMR spectra confirming CTP-phosphonoformate nucleotidyltransferase activity of His-SUMO-phpF. **A** Assay reaction mixture containing 50 mM HEPES (pH 7.25), 10 mM $MgCl_2$, 1 mM CTP, and 1 mM PF. **B** Assay reaction mixture after addition of 5 μg His-SUMO-phpF and incubation at 37°C for 1 h. **C** Assay reaction mixture after addition of 5 μg His-SUMO-phpF and 5 U PPase and incubation at 37°C for 1 h.

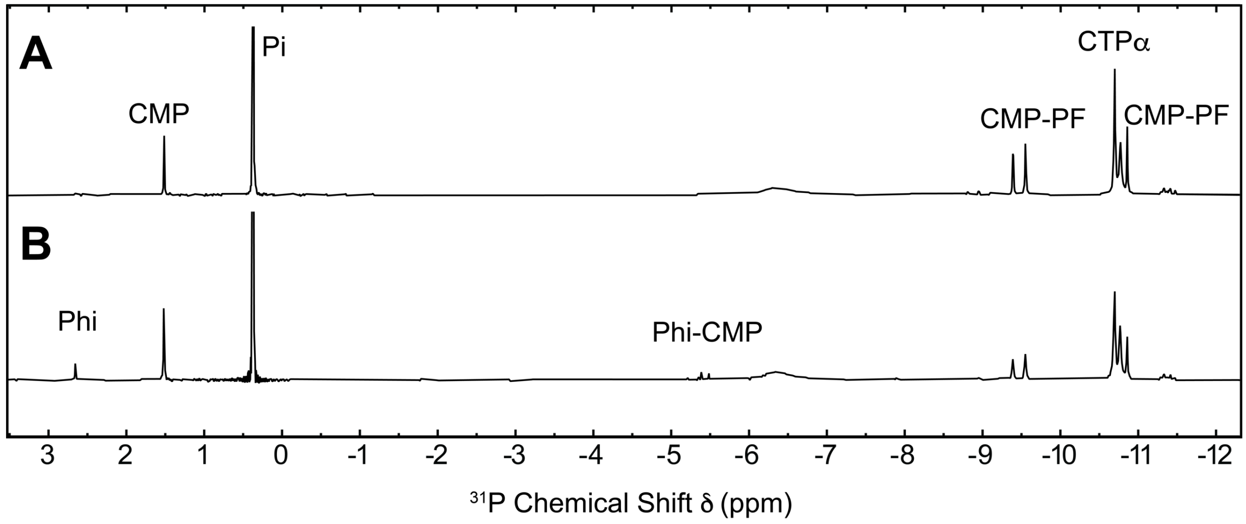


**Supplemental Figure S10.** Proton-decoupled $^{31}$P-NMR spectra demonstrating Phi production in trypsin-digested His-SUMO-phpF reaction mixtures. **A** Reaction mixture containing 50 mM MES (pH 6), 10 mM $MgCl_2$, 5 mM PF, 15 mM CTP, 5 μg His-SUMO-phpF, and 2.5 U PPase incubated for 2 h before addition of 1 μL trypsin. **B** Trypsin-digested sample after 24 h incubation at 37°C.

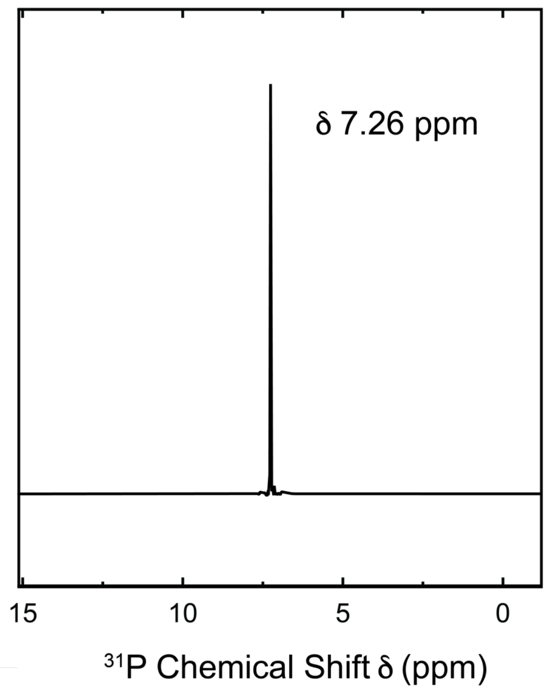


**Supplemental Figure S11.** Proton-decoupled $^{31}$P-NMR spectra of aminotris(methylenephosphonic acid) (ATMP) standard.
